# Identification of Brain Expression Quantitative Trait Loci Associated with Schizophrenia and Affective Disorders in Normal Brain Tissue

**DOI:** 10.1101/068007

**Authors:** Oneil G. Bhalala, Artika P. Nath, UK Brain Expression Consortium, Michael Inouye, Christopher R. Sibley

## Abstract

Schizophrenia and the affective disorders, here comprising bipolar disorder and major depressive disorder, are psychiatric illnesses that lead to significant morbidity and mortality worldwide. Whilst understanding of their pathobiology remains limited, large case-control studies have recently identified single nucleotide polymorphisms (SNPs) associated with these disorders. However, discerning the functional effects of these SNPs has been difficult as the associated causal genes are unknown. Here we evaluated whether schizophrenia and affective disorder associated-SNPs are correlated with gene expression within human brain tissue. Specifically, to identify expression quantitative trait loci (eQTLs), we leveraged disorder-associated SNPs identified from six Psychiatric Genomics Consortium and CONVERGE Consortium studies with gene expression levels in post-mortem, neurologically-normal tissue from two independent human brain tissue expression datasets (UK Brain Expression Consortium (UKBEC) and Genotype-Tissue Expression (GTEx)). We identified 6 188 and 16 720 cis-acting SNPs exceeding genome-wide significance (p<5x10^−8^) in the UKBEC and GTEx datasets, respectively. 1 288 cis-eQTLs were significant in a metaanalysis leveraging overlapping brain regions and were associated with expression of 15 genes, including three non-coding RNAs. One cis-eQTL, rs 16969968, results in a functionally disruptive missense mutation in *CHRNA5*, a schizophrenia-implicated gene. Meta-analysis identified 297 *trans*-eQTLs associated with 24 genes that were significant in a region-specific manner. Importantly, comparing across tissues, we find that blood eQTLs largely do not capture brain cis-eQTLs. This study identifies putatively causal genes whose expression in region-specific brain tissue may contribute to the risk of schizophrenia and affective disorders.

## Introduction

Schizophrenia and affective disorders, comprising bipolar disorder and major depressive disorder, constitute a significant global burden of disease. Worldwide it is estimated that more than 21 million individuals are living with schizophrenia, 60 million with bipolar disorder and over 400 million with major depressive disorder^1^. The consequences are staggering as these three diseases, which usually emerge in early-adulthood, accounted for over 90 million disability-adjusted life years in 2010^2^.

Ineffective management, which contributes to the enormous disease burden, is largely due to our lack of understanding about the pathobiology underlying these disorders. Family and twin studies have estimated heritability to be between 70% and 80% for schizophrenia and bipolar disorder and up to 40% for major depressive disorder^3^. This has prompted the establishment of genome-wide association studies (GWAS) to identify genetic variants associated with these disorders. Recent and large-scale GWAS organized by the Psychiatric Genomics Consortium (PGC) and CONVERGE Consortium have been published for schizophrenia^4^, bipolar disorder^5, 6^, major depressive disorder^7, 8^ as well as for a multiple-disorder analysis^9^. Results from these studies have suggested that schizophrenia, bipolar disorder and major depressive disorder may share common genetic architecture^10, 11^.

While GWAS have identified numerous loci associated with human diseases^12, 13^, understanding their roles in disease mechanism remains limited. Expression quantitative trait loci (eQTLs) are genetic variants that affect gene expression levels and may offer insights into mechanisms contributing to health and disease^14–16^. eQTLs can act in *cis*, meaning that the variant is associated expression of a gene within 1Mb, or in *trans*, with the variant located outside of this window. These two categories of eQTLs highlight the different avenues by which eQTLs may be exerting their influence in cellular processes^17^. Studies have leveraged GWAS data, particularly that of single nucleotide polymorphisms (SNPs), to identify eQTLs^18, 19^ Moreover, it is becoming evident that disease-associated variants are enriched for eQTLs^9, 20^. Hence, eQTL studies based on disease-based GWAS may offer insights into important molecular processes underlying pathobiology.

An important consideration in eQTL analyses is the source of tissue used for gene expression data. While identification of eQTLs is increased by studying associations in multiple tissues^21, 22^, other studies have found that analyses in disease-related tissues are enriched for disease-associated eQTLs^23, 24^, especially *trans^25–21^*. Recent availability of datasets detailing gene expression in various regions of human brain have now allowed for eQTL analyses in nervous tissue^28–30^.

In this study, we sought to identify eQTLs associated with schizophrenia and affective disorders in neurologically-normal post-mortem brain tissue. By leveraging gene expression data from UK Brain Expression Consortium (UKBEC)^29^ and Genotype-Tissue Expression (GTEx)^30^ with SNP data available from PGC, we identified cis-eQTLs that were pervasive across various brain regions. In contrast, trans-eQTLs were only detected in a region-specific manner. Results reported here may help prioritize future studies of GWAS SNPs associated with these disorders.

## Materials and Methods

### Collation of analysis-SNPs from GWAS data

Publically available GWAS data from six recent studies related to schizophrenia, bipolar disorder and major depressive disorder were included in this analysis (Figure 1). Data were downloaded from the PGC website (https://www.med.unc.edu/pgc). SNPs were collated from the following studies: PGC-SCZ2^4^ for schizophrenia, PGC-BIP^5^ and PGC-MooDs^6^ for bipolar disorder, PGC-MDD^7^ and CONVERGE^8^ for major depressive disorder, and PGC-Cross Disorder Analysis^9^ for multiple disorders (schizophrenia, bipolar disorder, major depressive disorder, autism spectrum disorder and attention-deficit hyperactivity disorder). Of note, PGC-MooDs analysis included samples from the PGC-BIP study; PGC-Cross Disorders analysis included samples from the PGC-SCZ2, PGC-BIP and PGC-MDD studies. We included overlapping studies to maximize the number of disorder-associated SNPs in our analysis as there may be loci identified in one study but not in another.

**Figure 1.**
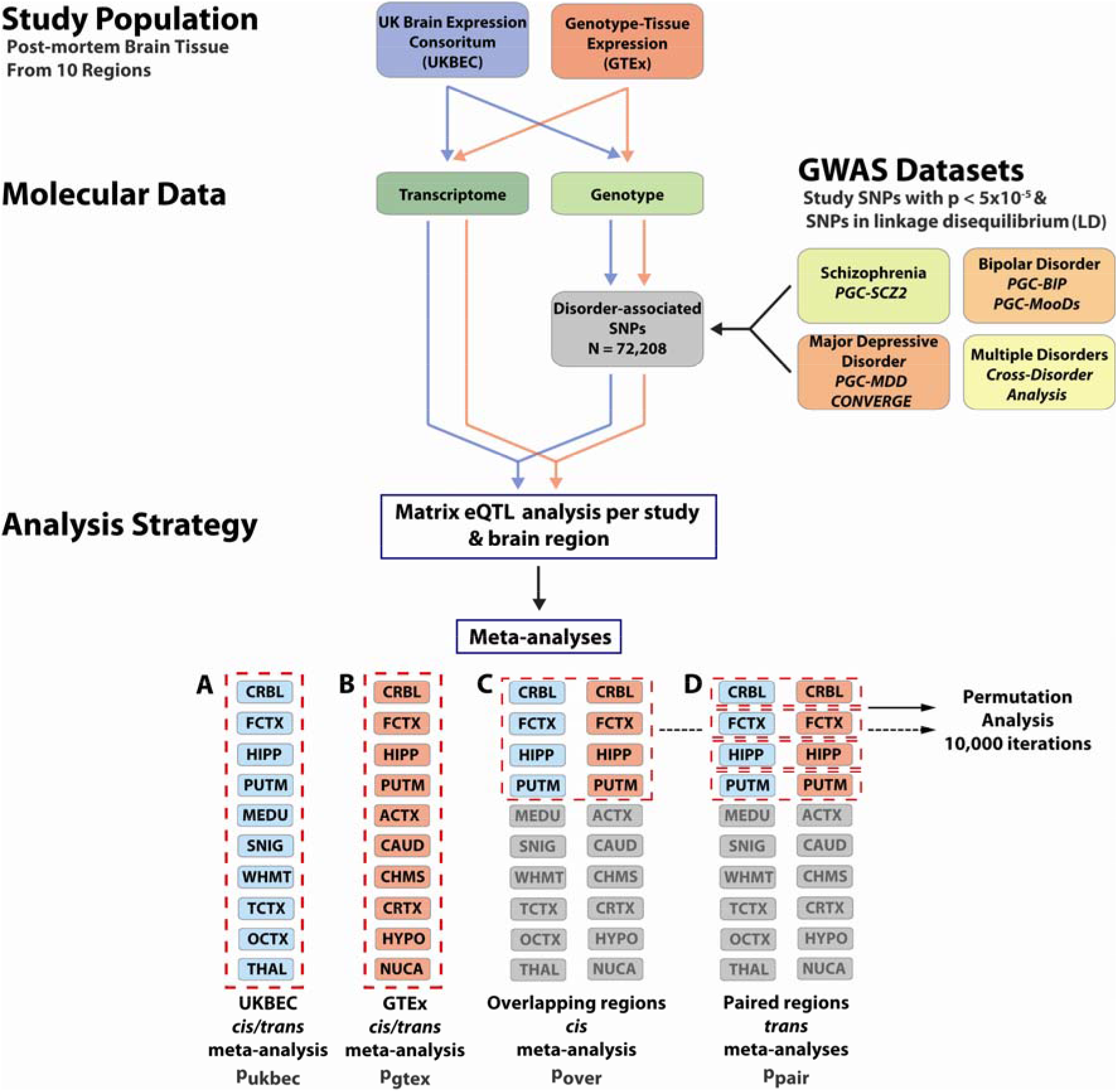
Study design for identification of brain eQTLs. Two studies with genotype and transcriptomic data (UKBEC and GTEx) were analysed for eQTLs based on SNPs that were found to be associated with schizophrenia, bipolar disorder and major depressive disorder through six GWAS studies. UKBEC consisted of samples from neurologically-normal individuals; samples from individuals with neurological disease (see Methods) were removed from GTEx prior to analyses. For each study, each of the ten brain regions was analysed separately by Matrix eQTL for identification of *cis-* and trans-eQTLs. Matrix eQTL results were meta-analysed in the following manner: *A,B*, within each study, all ten regions were meta-analysed for both *cis-* and trans-eQTLs (p-values are denoted as puk_bec_ and p_gtex_, respectively); *C*, overlapping regions between the two studies were meta-analysed for *cis*-eQTLs (p_over_); and *D*, each pair of overlapping regions was meta-analysed separately for trans-eQTLs (p_pa_ir). *CRBL*, cerebellum; *FCTX*, frontal cortex; *HIPP*, hippocampus; *PUTM*, putamen; *MEDU*, medulla; *SNIG*, substantia nigra; *WHMT*, white matter; *TCTX*, temporal cortex; *OCTX* occipital cortex; *THAL*, thalamus; *ACTX* anterior cingulate cortex; *CAUD*, caudate; *CHMS*, cerebellar hemisphere; *CRTX*, cortex; *HYPO*, hypothalamus; *NUCA*, nucleus accumbens.

SNPs with a study p-value < 5 x 10^−5^ were included in the analysis. For SNPs from all studies except PGC-SCZ2, we also obtained SNPs that are in moderate-high linkage disequilibrium (LD, R^2^ ≥ 0.5) with the study-SNPs^31^ using the web-based tool rAggr (http://raggr.usc.edu/). LD-analysis settings were as follows: CEU population from 1000 Genomes Phase 3 October 2014 release, build hg19, minimum minor allele frequency = 0.001, maximum distance = 500kb, maximum Mendelian errors = 1, Hardy-Weinberg p-value cutoff = 0, and minimum genotype = 75%. Study-SNPs from PGC-BIP, PGC-MDD and PGC-Cross Disorder Analysis were lifted from hg18 to hg19 prior to LD-analysis. PGC-SCZ2 had nearly 46,000 study SNPs with p < 5 x 10^−5^, so additional SNPs in LD with these were not obtained. Supplementary Table 1 indicates the number of study-SNPs and analysis-SNPs (study-SNPs and those SNPs in LD with them) per study.

### Brain expression and genotype data

Genotype and gene expression data were obtained from the UK Brain Expression Consortium (UKBEC)^29^. These data contained samples from 134 neuropathologically-free individuals from the following brain regions: cerebellum (n=96), frontal cortex (n=97), hippocampus (n=96), medulla (n=95), occipital cortex (n=98), putamen (n=99), substantia nigra (n=73), temporal cortex (n=86), thalamus (n=95), and white matter (n=97). Processing of genotype is previously described^29^. Briefly, these data included ~5.88 million imputed (1000 Genome, March 2012 release) and typed SNPs. Raw expression data from Affymetrix Human Exon 1. 0 ST microarrays were processed as described previously with minor modifications. Specifically, all ~5 million probes were initially re-mapped to Ensembl v75 annotations using Biomart. Only probe sets containing three or more probes free of the ‘polymorphism-in-probe’ problem were used for subsequent analysis^32^. Following RMA normalisation and background filtering, we calculated gene-level estimates by taking the winsorised mean of all probe sets for a given gene. Prior to eQTL detection, these were covariate-corrected for sex, cause of death, post-mortem interval and RIN.

Data from Genotype-Tissue Expression (GTEx)^30^ were also included in our study. The following brain regions were analysed: anterior cingulate cortex (BA24) (n=72), caudate (n=100), cerebellar hemisphere (n=89), cerebellum (n=103), cortex (n=96), frontal cortex (BA9) (n=92), hippocampus (n=81), hypothalamus (n=81), nucleus accumbens (n=93), and putamen (n=82). Whole blood tissue (n=338) was also analysed, serving as a comparison to brain tissues. Genotype and expression processing is described at http://gtexportal.org. Briefly, dataset contained ~11.55 million imputed (1000 Genomes, August 2012 release) and typed autosomal variants and ~26,000 transcripts (RPKM data from RNA-SeQC with quantile normalization). To make this dataset similar to UKBEC, we excluded individuals from GTEx that were identified as having neurological diseases based on description of their comorbidities/cause of death or variables (variables excluded were: MHALS (ALS), MHALZDMT (Alzheimer’s or dementia), MHALZHMR (Alzheimer’s), MHPRKNSN (Parkinson’s disease), MHDMNTIA (dementia with unknown cause), MHDPRSSN (major depression), MHSCHZ (schizophrenia), MHENCEPHA (active encephalitis), MHJAKOB (Cruetzfeldt Jakob relatives), MHMENINA (active meningitis), MHMS (multiple sclerosis), MHSZRSU (unexplained seizures)).

### eQTL analysis

The R package Matrix eQTL^33^ was used to identify eQTLs. Cis-eQTLs were defined as SNPs within 1Mb of the transcription start site; trans-eQTLs were those SNPs outside this region. Study genotype data was limited to those that were present in the set of analysis-SNPs described above. Due to the relatively low sample numbers, there is a possibility that an eQTL signal may be deemed significant if it is showing a very strong effect in only a few individuals. To minimize this event, analysis-SNPs that had a minor allele frequency < 5% within the analysis population were excluded. For UKBEC, gene expression input was based on residuals from linear regression of gene expression and genotype with the following covariates: sex, cause of death, age at death, post-mortem interval and RIN. Hence, no covariate data were separately considered in Matrix eQTL. Covariate data consisting of three genotyping principal components (PCs), array platform, gender and PEER factors were utilised for GTEx analysis. eQTL analysis, for both *cis* and *trans*, was performed independently for each region per study. eQTL (beta) effect sizes are given as standardized expression units (EU) per allele.

### Meta-analyses across brain regions and studies

To determine which eQTLs were significant in multiple regions/studies, we performed a meta-analysis of test statistics (eQTL effect size estimates and standard errors) calculated by Matrix eQTL using the metagen function in the R package meta with a random-effects model. False discovery rate (FDR) was calculated using R p.adjust with the Benjamini-Hochberg method. Unless otherwise stated we used a genome-wide significant p-value of p < 5 x 10^−8^ as our cutoff for calling a significant eQTL. For completeness, in supplementary tables, we additionally report the number of eQTLs passing a FDR threshold of FDR < 0.05. Four groups of meta-analyses were performed (Figure 1): A) of all ten regions in UKBEC for *cis* and *trans*, separately (p-valued denoted as p_ukbec_); B) of all ten regions in GTEx for *cis* and *trans*, separately (p_gtex_); C) of four overlapping regions (cerebellum, frontal cortex, hippocampus and putamen) between UKBEC and GTEx for *cis* (p_over_); and D) by pairs of overlapping regions (separate meta-analysis for each cerebellum, frontal cortex, hippocampus and putamen) for *trans* (ppair). To increase power for analyses C and D, only eQTLs that had the same effect direction in each study per region were analysed. The first ten principal components of the UKBEC-GTEx meta-analysis for each of the overlapping regions are shown in Supplementary Figures 1–4.

### Permutation analysis to test meta-analysis results

Permutation analysis was used to test the robustness of eQTLs identified in the meta-analysis of overlapping regions in UKBEC and GTEx datasets (analyses C and D above, Figure 1). 10 000 iterations were performed for each region-study combination by shuffling sample IDs in the gene expression file and using Matrix eQTL as described above. Meta-analysis of each permuted iteration per combination yielded a meta p-value. Permuted p-value was calculated as the number of meta p-values equal to or less than the nominal p-value divided by the number of iterations (10 000). Those eQTLs with permuted p-values less than 0.05 were considered significant.

## Results

### Identification of disease-associated eQTLs in UKBEC and GTEx

We sought to determine if SNPs associated with schizophrenia and affective disorders also served as eQTLs in brain tissue. Study-SNPs identified in six GWAS were included in analysis: PGC-SCZ2^4^ for schizophrenia, PGC-BIP^5^ and PGC-MooDs^6^ for bipolar disorder, PGC-MDD^7^ and CONVERGE^8^ for major depressive disorder, and PGC-Cross Disorder^9^ which included schizophrenia, bipolar disorder, major depressive disorder, autism spectrum disorder and attention-deficit hyperactivity disorder (Figure 1). In order to maximise detection of eQTLs^31^, we included SNPs that are in moderate to high LD (R^2^ >= 0.5) with study-SNPs using rAggr (Supplementary Table 1). Combining study-SNPs and those in LD with them yielded 72 208 analysis-SNPs across the six GWAS.

Genotype and gene expression data were obtained from UKBEC^29^ and GTEx^30^, which contained information of post-mortem human tissue from samples independent from each other and from the GWAS used to identify analysis-SNPs. Of the 72 208 analysis-SNPs, 58 325 and 58 133 were present in both UKBEC and GTEx, respectively. Expression data originated from ten UKBEC regions (cerebellum, frontal cortex, hippocampus, putamen, temporal cortex, occipital cortex, substantia nigra, white matter, thalamus, medulla) and ten GTEx regions (anterior cingulate cortex (BA24), caudate, cerebellar hemisphere, cerebellum, cortex, frontal cortex (BA9), hippocampus, hypothalamus, nucleus accumbens, putamen). Samples in UKBEC were neurologically-disease free. Those samples with neurological disease in GTEx data were removed from analysis (see Methods), allowing for identification of eQTLs in disease-free tissue.

To identify cis-eQTLs in these tissues, we utilised an additive linear model with Matrix eQTL^33^. Using this methodology, we initially analyzed cis-eQTLs in each of the ten regions within UKBEC and GTEx datasets, separately (Figure 1, analyses A,B, Supplementary Figures 5,6). The number of detected cis-eQTLs using a stringent threshold of p < 5 x 10^−8^ varied considerably between studies and regions (Supplementary Table 2). For example, in the UKBEC dataset, the cerebellum harboured 1 234 unique cis-eQTLs, while none were detected in substantia nigra or medulla. Within the GTEx dataset, 2 879 and 696 were detected in the cerebellum and anterior cingulate cortex, respectively. We meta-analysed *cis*-eQTLs across all ten regions in UKBEC and GTEx separately (Supplementary Figure 7, Supplementary Tables 3 and 4). When collectively considered, 6 188 and 16 720 unique *cis*-eQTLs were significant (pukbec/gtex < 5 x 10^−8^) in the UKBEC and GTEx data, respectively. These cis-eQTLs were associated with expression variation of 146 and 506 genes, respectively. 3 169 cis-eQTLs, associated with 46 genes, were shared between these datasets at a threshold of p_u_k_bec/gtex_ < 5 x 10^−8^ (Supplementary Figure 8).

The number of *trans*-eQTLs that were found to be significantly associated in any of the individual ten regions per study was greatly reduced compared to cis-eQTLs. Zero *trans-* eQTLs were detected in meta-analysis of all ten UKBEC regions (Figure 1, analysis A). In the GTEx dataset (Figure 1, analysis B), 103 trans-eQTLs were detected and associated with expression of two non-coding RNAs of unknown function located on chromosome 8: *RNF5P1* (most significant SNP is rs3130349 on chromosome 6p21.32, p_gtex_ = 8.3 x 10^−222^) and *FAM66A* (most significant SNP is rs7832708 on chromosome 8p23.1, p_gtex_ = 8.8 x 10^−62^).

### Meta-analysis of eQTLs in overlapping brain regions across studies

To generate a high-confidence list of eQTLs shared between datasets as well as to discover additional ones, we next sought to determine if eQTLs identified in either UKBEC or GTEx data were significant when assessed across both studies. Four brain regions overlapped between the two datasets: cerebellum, frontal cortex, hippocampus and putamen. Test-statistics restricted to these regions were meta-analysed (Figure 1, analyses C and D). This reduced the aforementioned 3 169 overlapping cis-eQTLs to 1 288 cis-eQTLs that were significant with pover <5 x 10^−8^ (Supplementary Table 5). To further validate cis-eQTL associations, we performed permutation analysis of 10 000 iterations for each expression dataset and brain region (resulting in eight expression-region groups meta-analysed 10 000 times each). Each of the 1 288 cis-eQTLs with p_over_ < 5 x 10^−8^ had a permuted p-value < 1 x 10^−4^.

The 1 288 cis-eQTLs were associated with expression of 15 genes (Figure 2A, Table 1). Of these, 784 cis-eQTLS associated with 10 genes also had p_ukbec_ < 5 x 10^−8^ in the UKBEC metaanalysis of the four overlapping brain regions and 714 cis-eQTLs associated with 6 genes also had pgt_ex_ < 5 x 10^−8^ in the GTEx meta-analysis of the four overlapping brain regions. Demonstrating reproducibility between the two studies, 568 of the cis-eQTLs within these two groups had both pukbec < 5 x 10^−8^ and pgtex < 5 x 10^−8^ (Figure 2B). Moreover, an additional 358 cis-eQTLs were detected when both UKBEC and GTEx datasets were leveraged in combination, thus demonstrating the benefit of a combined analysis. Mean and median of the (absolute value of) effect sizes (irrespective of p_over_) were 0.33 and 0.34 EU per allele, respectively (Supplementary Figure 9). The largest effect size was observed with rs9461434 (-0.70 ± 0.12 EU per allele for *AA* relative to GG), associated with the expression of the pseudogene *ZNF603P* (p_over_ = 2.1 x 10^−8^, FDR = 1.4 x 10^−6^, Supplementary Figure 10).

**Figure 2.**
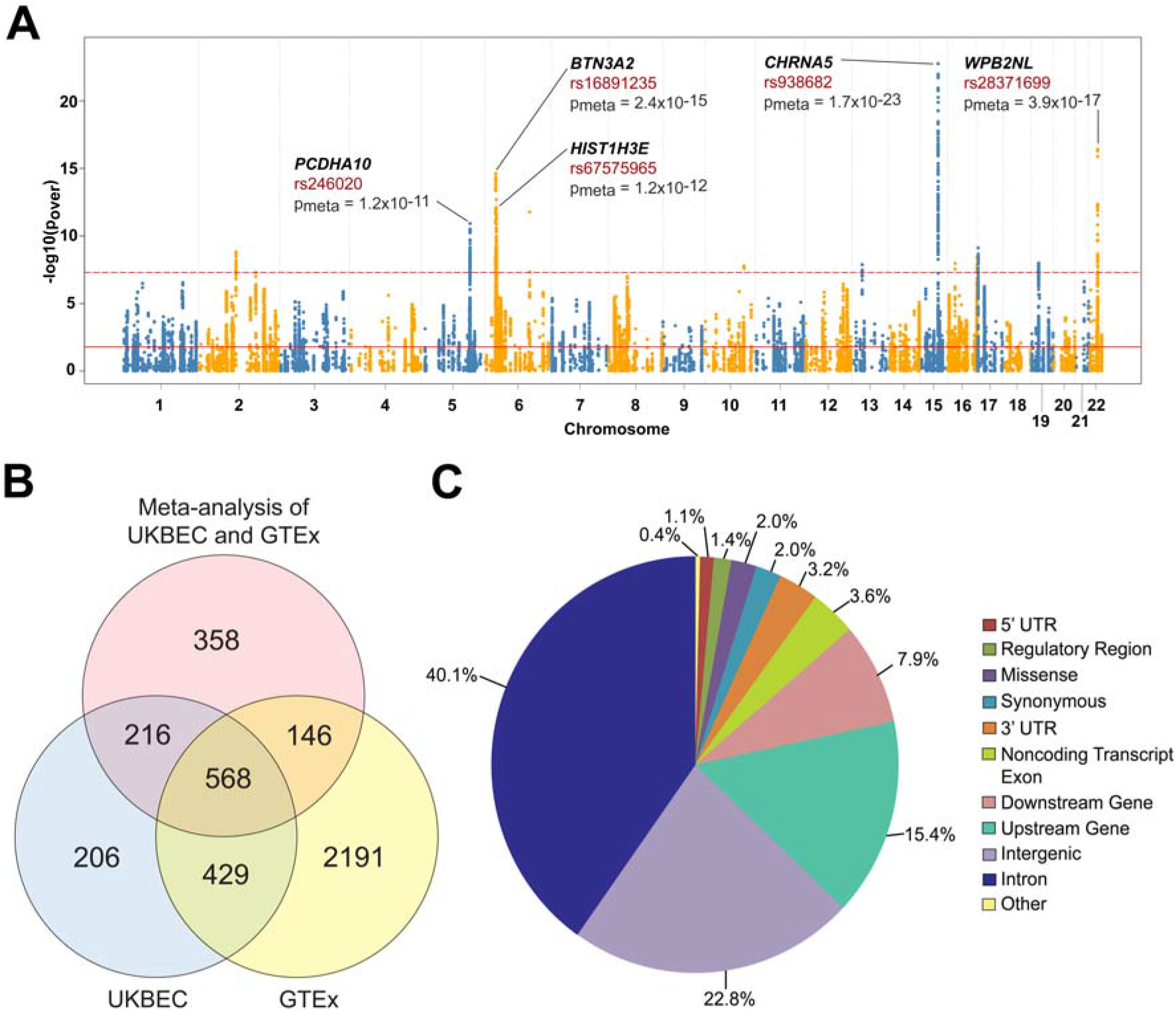
Manhattan plot and effect predictions of meta-analysed *cis*-eQTLs. *A*, −log10(p_over_) are plotted for cis-eQTLs identified in a meta-analysis of four overlapping brain regions between UKBEC and GTEX (Figure 1, analysis C). Top five protein-coding genes associated with these eQTLs are provided, along with the associated cis-eQTL and pover. Solid red line denotes threshold at which FDR = 0.05. Dashed red line denotes threshold at which p-value = 5 x 10^−8^. *B*, Overlap of cis-eQTLs between meta-analysis of four brain regions common between UKBEC and GTEx (Figure 1, analysis C) with p_over_ < 5 x -10^−8^, meta-analysis of those regions in UKBEC alone with pukbec < 5 x 10^−8^ and meta-analysis of those regions in GTEx alone with pgtex < 5 x 10^−8^. C, SNP effect prediction (from Ensembl Variant Effect Predictor) for 1 288 cis-eQTLs with p_over_ < 5 x 10^−8^. Overlapping brain regions are cerebellum, frontal cortex, hippocampus and putamen.

**Table 1:** Most significant c/s-eQTL for each associated gene.

| Gene | Gene Function | eQTL | Alleles<sup>†</sup> | eQTL Location | $\beta_{\text{over}} \pm \text{se}$ | $p_{\text{over}}$ | FDR |
| --- | --- | --- | --- | --- | --- | --- | --- |
| ***CHRNA5*** | Ligand-gated ion channel | rs938682 | A/G | Chr 15, *CHRNA3* intron | $0.26 \pm 0.03$ | 1.7E-23 | 5.8E-19 |
| ***WBP2NL*** | Sperm-specific WW domain-binding protein | rs28371699 | A/C | Chr 22, *CYP2D6* intron | $-0.13 \pm 0.02$ | 3.9E-17 | 6.6E-14 |
| ***BTN3A2*** | Immunoglobulin superfamily | rs16891235 | T/C | Chr 6, *HIST1H1A* exon (missense variant) | $0.15 \pm 0.02$ | 2.4E-15 | 2.6E-12 |
| ***ZNF603P*** | Non-coding RNA | rs200976 | A/G | Chr 6, intergenic variant | $-0.48 \pm 0.06$ | 2.0E-13 | 1.4E-10 |
| ***HIST1H3E*** | Nucleosome component | rs67575965 | A/G | Chr 6, intergenic variant | $-0.30 \pm 0.04$ | 1.2E-12 | 5.9E-10 |
| ***C6orf3*** | Non-coding RNA | rs6912446 | T/A | Chr 6, *TRAF3IP2* intron | $0.17 \pm 0.02$ | 1.7E-12 | 7.9E-10 |
| ***PCDHA10*** | Calcium-dependent cell-adhesion | rs246020 | A/G | Chr 5, intron in multiple genes including *PCDHA10* | $-0.35 \pm 0.05$ | 1.2E-11 | 4.2E-09 |
| ***SRR*** | Catalyzes the synthesis of D-serine | rs395566 | C/T | Chr 17, *SMG6* intron | $-0.16 \pm 0.03$ | 7.6E-10 | 1.5E-07 |
| ***SEPT10*** | Filament-forming cytoskeletal GTPase | rs6752531 | G/A | Chr 2, *SEPT10* intron | $0.12 \pm 0.02$ | 1.5E-09 | 2.7E-07 |
| ***GAS8*** | Cytoskeletal linker | rs11645893 | G/C | Chr 16, *FANCA* intron | $0.08 \pm 0.01$ | 4.1E-09 | 5.5E-07 |
| ***MAU2*** | Association of cohesion complex with chromatin during interphase | rs8101499 | G/A | Chr 19, downstream of *MAU2* | $0.06 \pm 0.01$ | 1.0E-08 | 9.3E-07 |
| ***RRN3*** | Transcription initiation | rs11075255 | G/A | Chr 16, intron in *RRN3* and *PDXDC1* | $-0.12 \pm 0.02$ | 1.1E-08 | 9.4E-07 |
| ***SMIM2-AS1*** | Non-coding RNA | rs12869383 | A/G | Chr 13, intergenic | $-0.52 \pm 0.09$ | 1.3E-08 | 9.8E-07 |
| ***GSTO2*** | Glutathione-dependent thiol transferase activity | rs7079338 | C/T | Chr 10, *GSTO2* intron | $0.32 \pm 0.06$ | 1.6E-08 | 1.2E-06 |
| ***ZSCAN31*** | Transcription factor | rs2142730 | T/C | Chr 6, *PGBD1* intron | $-0.34 \pm 0.06$ | 2.9E-08 | 1.7E-06 |

<sup>†</sup>First allele listed is the reference allele for effect size. For example, compared to G, A has a $\beta$ of $0.26 \pm 0.03$ EU per allele with respect to expression levels of *CHRNA5*.

Of the 1 288 cis-eQTLs with p_over_ < 5 x 10^−8^, 40.1% were located in introns and 22.8% within intergenic regions (Figure 2C), as classified by Ensembl Variant Effect Predictor. The most significant cis-eQTL was rs938682 on chromosome 15q25.1, correlated with expression of *CHRNA5* (p_over_ = 1.7 x 10^−23^, FDR = 5.8 x 10^−19^), which encodes the α5 subunit of the nicotinic cholinergic receptor. This eQTL has an effect size of 0.26 ± 0.03 EU per allele for *AA* (allele frequency of *A =* 0.79 in 1000 Genomes CEU population) relative to *GG* (Figure 3A,B). Interestingly, the eQTL is located within an intron of *CHRNA3*, whose protein interacts with *CHRNA5* to form a nicotinic acetylcholine receptor^34^. There was no correlation between *CHRNA3* and *CHRNA5* expression with a Pearson R^2^ = 0.02. Moreover, no significant cis-eQTLs associated with *CHRNA3* expression were observed.

**Figure 3.**
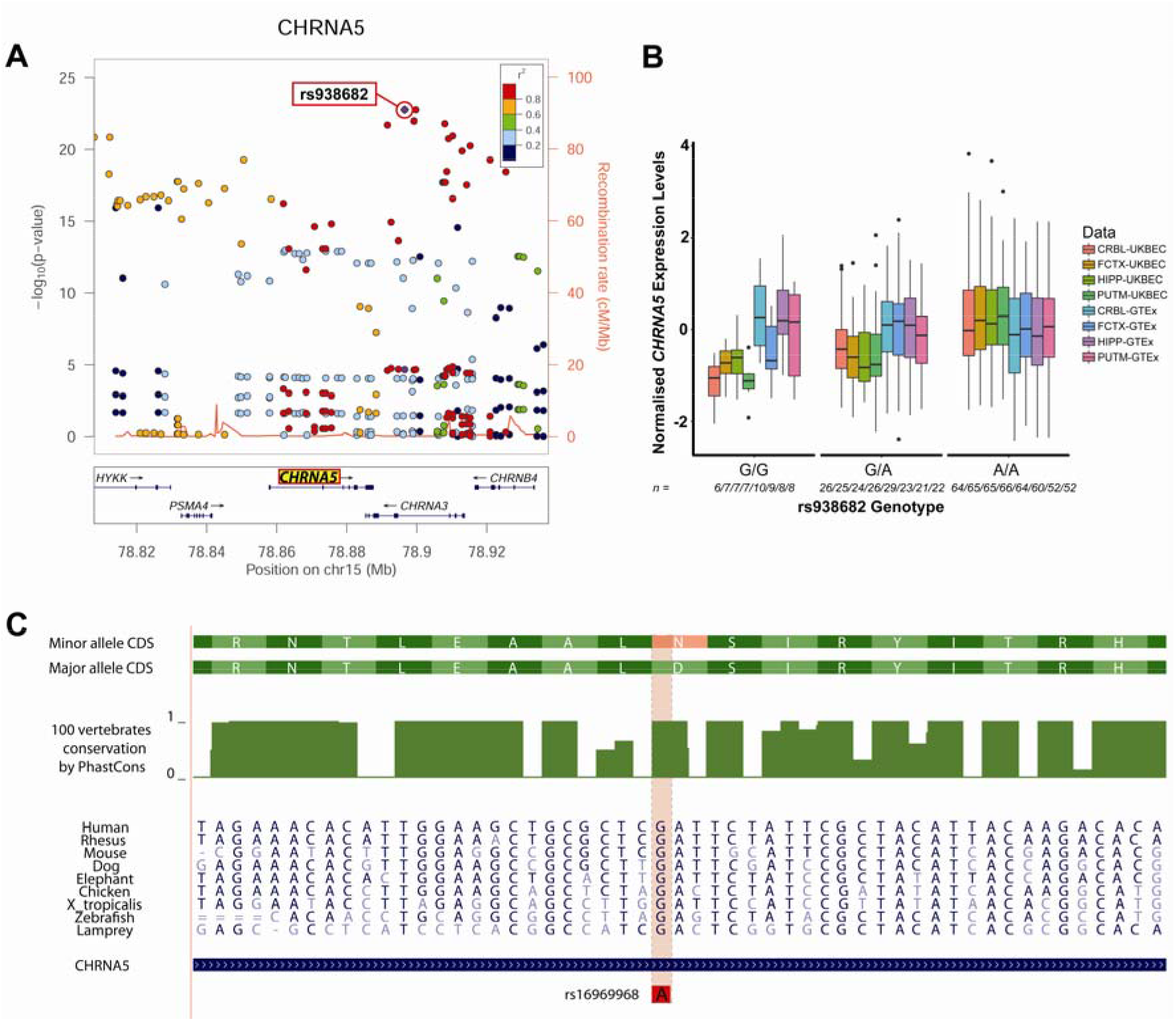
Mapping and gene expression effects of *CHRNA5* associated eQTLs. *A*, LocusZoom plot for the most significant cis-eQTL (rs938682) identified in meta-analysis of UKBEC and GTEx (Figure 1, analysis C, Table 1). This SNP is *cis* to *CHRNA5*, denoted in yellow highlight. Other SNPs that are within the study and *cis* to *CHRNA5* are plotted for their pover (left y-axis) and LD-value (r^2^ denoted by color of circle, calculated as relative to rs938682). Recombination rate of genomic region is plotted in red. *B*, Boxplot of gene expression levels (normalised separately for UKBEC and GTEx per region) by genotype of the rs938682 eQTL. Vertical lines for each plot captures data between −1.5 x interquartile rage and 1.5 x interquartile range, with outliers depicted as black circles. The number of individuals with a particular genotype per study-region is denoted in italics. *C*, Amino acid conservation plot for CHRNA5. Missense mutation resulting from the eQTL rs16969968 is highlighted.

Interestingly, 26 cis-eQTLs (Supplementary Table 6) were classified as leading to missense mutations. Five eQTLs resulted in missense mutations in genes whose expression with which they were associated. The most significant of these was rs16969968 (p_over_ = 8.4 x 10^−13^), which is located within exon 5 of *CHRNA5*, a highly conserved region (Figure 3C). This finding lends further support to the involvement of *CHNRA5* in these diseases. The minor allele *A* encodes for an amino acid change to asparganine from aspartic acid (major allele *G*) at position 398, which may affect receptor function (see Discussion). The effect size of GG, relative to AA, is 0.23 ± 0.03 EU per allele (Supplementary Figure 11), thus implying that both expression changes and receptor activity could be contributing to the effect of this variant.

As noted above, only two trans-eQTLs were detected in GTEx. As expected, neither remained significant when the four overlapping regions were meta-analysed together. However, meta-analysis of each region across both studies identified varying amount of trans-eQTLs with p_pair_ < 5 x 10^−8^ (Figure 1, analysis D; Table 2; Supplementary Tables 7-10). Twelve trans-eQTLs associated with five genes within the cerebellum; 255 trans-eQTLs associated with twelve genes within the frontal cortex; eleven trans-eQTLs associated with two genes within the hippocampus; and 19 trans-eQTLs associated with five genes within the putamen (Figure 4^35^). These trans-eQTLs remained significant after permutation testing, with each having a permuted p-value < 1 x 10^−4^. In the frontal cortex, nearly 70% of trans-eQTLs were located within the *MHC* locus on chromosome 6p21 and were associated with expression of *DDX17* on chromosome 22q13.1, which encodes for a RNA-binding protein. While the specific role of this protein, also known as p72, in brain is still uncertain, it is involved in numerous aspects of RNA metabolism, including promoting microRNA biogenesis and reducing viral RNA stability^36, 37^ *DDX17* is also a binding-partner to the transcription factor SOX2^38^, thereby potentially influencing master regulation of gene expression.

**Figure 4.**
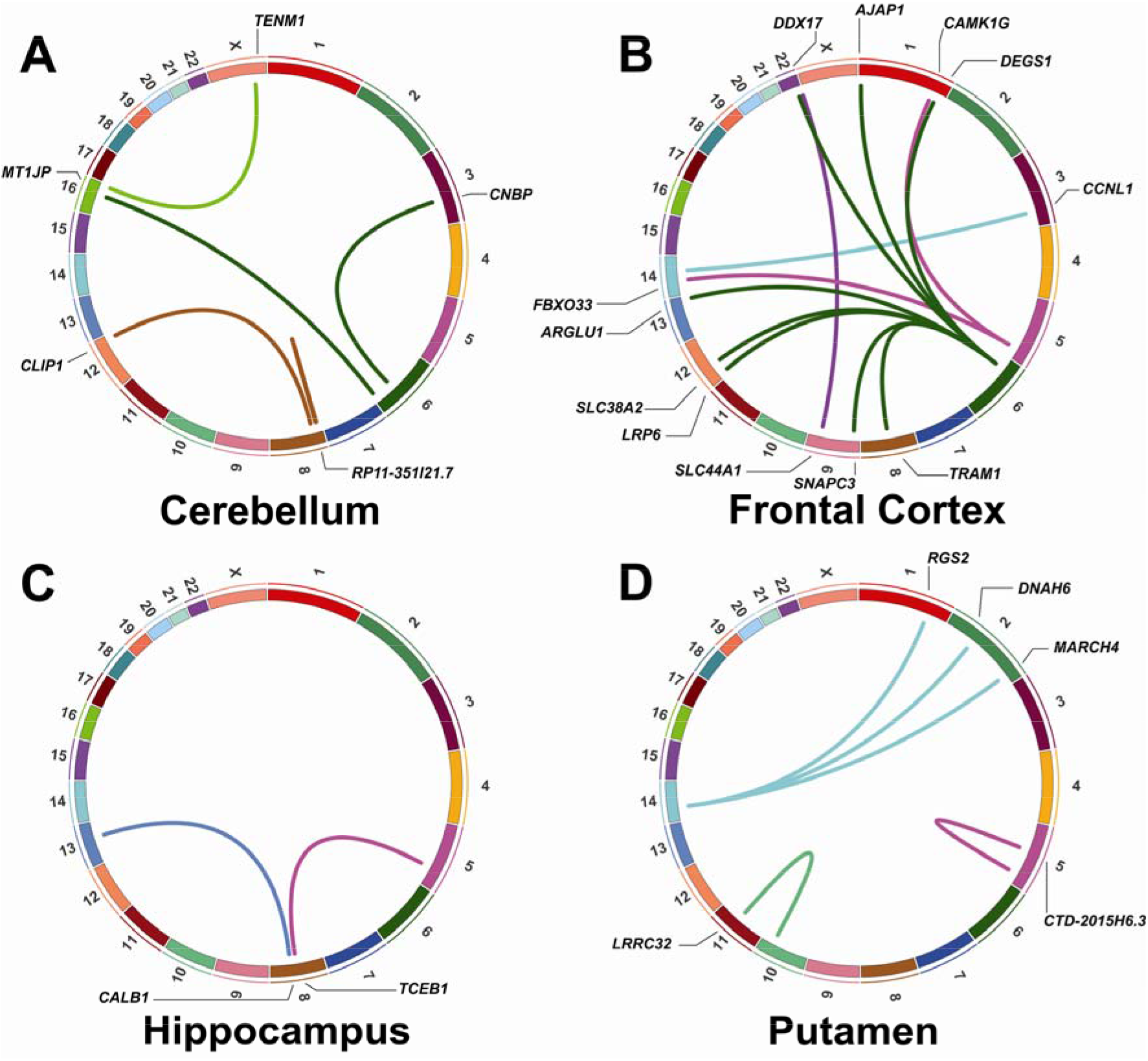
Circos^35^ plots of *trans*-eQTLs from meta-analysis by brain region. Locations of trans-eQTLs (with p_pair_ < 5 x 10^−8^ per overlapping brain region, *A-D*) and their respective associated genes. Chromosomal location of the eQTL is denoted by the color of the connecting line. For example, within the putamen *(D)*, trans-eQTLs are located on chromosome 14 (light blue) and are associated with expression of three genes on chromosomes 1 and 2.

**Table 2:** Most significant *trans-eQTL* for each associated gene by brain region.

| | eQTL | eQTL Location | Gene | Gene Chromosome | $\beta_{\text{pair}} \pm \text{se}$ | $p_{\text{pair}}$ | FDR* |
| --- | --- | --- | --- | --- | --- | --- | --- |
| Cerebellum | rs1883137 | 6q21 | *CNBP* | 3 | 0.22 ± 0.04 | 6.7E-09 | 2.9E-06 |
| | rs1995942 | 6q27 | *MT1JP* | 16 | 0.35 ± 0.06 | 9.3E-09 | 2.9E-06 |
| | rs6601450 | 8p23.1 | *RP11-351I21.7* | 8 | -0.42 ± 0.07 | 2.2E-08 | 4.3E-06 |
| | rs7839435 | 8p21.1 | *CLIP1* | 12 | 0.21 ± 0.04 | 3.8E-08 | 4.3E-06 |
| | rs4843852 | 16q24.1 | *TENM1* | X | 0.25 ± 0.05 | 4.9E-08 | 4.9E-06 |
| Frontal Cortex | rs28895257 | 6p21.32 | *DDX17* | 22 | 0.21 ± 0.03 | 3.1E-14 | 2.6E-11 |
| | rs72795371 | 5q33.1 | *CAMK1G* | 1 | 0.40 ± 0.07 | 1.9E-09 | 1.7E-08 |
| | rs6876723 | 5q33.1 | *FBXO33* | 14 | 0.26 ± 0.05 | 7.1E-09 | 6.0E-08 |
| | rs522308 | 6p21.32 | *SLC38A2* | 12 | 0.24 ± 0.04 | 7.8E-09 | 6.6E-08 |
| | rs9268659 | 6p21.32 | *SNAPC3* | 9 | -0.24 ± 0.04 | 9.4E-09 | 7.7E-08 |
| | rs5018660 | 6p21.32 | *LRP6* | 12 | 0.20 ± 0.04 | 9.5E-09 | 7.8E-08 |
| | rs9271620 | 6p21.32 | *TRAM1* | 8 | 0.18 ± 0.03 | 1.5E-08 | 1.2E-07 |
| | rs9268458 | 6p21.32 | *AJAP1* | 1 | 0.20 ± 0.03 | 2.2E-08 | 1.6E-07 |
| | rs9271763 | 6p21.32 | *ARGLU1* | 13 | 0.21 ± 0.04 | 2.2E-08 | 1.6E-07 |
| | rs9271480 | 6p21.32 | *DEGS1* | 1 | 0.26 ± 0.05 | 2.7E-08 | 1.8E-07 |
| | rs7142068 | 14q23.3 | *CCNL1* | 3 | 0.37 ± 0.07 | 4.1E-08 | 2.5E-07 |
| | rs145069606 | 22q13.33 | *SLC44A1* | 9 | -0.48 ± 0.09 | 4.8E-08 | 2.5E-07 |
| Hip | rs7448129 | 5q23.3 | *TCEB1* | 8 | 0.43 ± 0.08 | 1.7E-08 | 2.7E-06 |
| | rs1831189 | 13q21.33 | *CALB1* | 8 | -0.35 ± 0.06 | 3.4E-08 | 3.0E-06 |
| Putamen | rs11167589 | 5q33.1 | *CTD-2015H6.3* | 5 | 0.42 ± 0.07 | 8.3E-10 | 6.3E-07 |
| | rs57275892 | 14q13.2 | *MARCH4* | 2 | -0.28 ± 0.05 | 1.8E-08 | 1.4E-06 |
| | rs11817486 | 10q24.31 | *LRRC32* | 11 | -0.36 ± 0.06 | 2.8E-08 | 1.8E-06 |
| | rs12887279 | 14q13.2 | *RGS2* | 1 | -0.29 ± 0.05 | 3.9E-08 | 1.9E-06 |
| | rs57725522 | 14q13.2 | *DNAH6* | 2 | -0.39 ± 0.07 | 4.2E-08 | 1.9E-06 |

\* FDR values were calculated per brain region. *Hip*, Hippocampus.

### Overlap with whole blood tissue eQTLs

Previous eQTL analyses for schizophrenia and affective disorders have relied on transcriptome data collected from more readily available biological specimens such as whole blood^39–41^ or single brain regions^41^. An important question is whether disease-related eQTLs can be detected in samples that are more accessible and available from living patients. If so, this may facilitate larger study designs in the future where high quality biological material is more readily selected. To test whether our significant cis-eQTLs detected brain regions can be identified in more clinically-accessible tissue, we assessed the associations of 1 288 *cis*-eQTLs with pover < 5 x 10^−8^ (Figure 1, analysis C) in whole blood tissue within the GTEx database (n = 282 after removal of individuals with neurological disease (see Methods)). Of these, 472 had a p_blood_ < 0.05. However, only 143 (11%) reached our stringent threshold and had a p_blood-GT_E_x_ < 5 x 10^−8^ (Supplementary Figure 12A). Looking in more detail, of the fifteen genes associated with brain cis-eQTLs from our cross-study meta-analysis, only twelve had detectable expression in the GTEx whole blood samples. Moreover, only *BTN3A2* was associated with blood cis-eQTLs having p_blood-GTEx_ < 5 x 10^−8^. The most significant brain *cis*-eQTL in whole blood was rs71557332 (chromosome 6p22.2), which correlated with *BTN3A2* expression (p_blood-GTEx_ = 1.8 x 10^−52^ and FDR = 3.6 x 10^−49^). Of note, the direction of the effect was the same as that seen in the brain eQTL; the effect size was 0.98 ± 0.05 EU per allele in blood compared to an effect size of 0.55 ± 0.10 EU per allele in brain (Supplementary Figure 12B,D). Contrastingly, the top brain cis-eQTL, rs938682, associated with *CHRNA5* expression, only had a p_blood-GTEx_ = 0.03 and FDR = 0.29 (effect direction was the same).

Similar results were observed when we tested for overlap of cis-eQTLs detected in the largest meta-analysis of peripheral whole blood to date (n = 5 311, no overlap with GTEx whole blood data)^42^. In this analysis, only 69 our of brain cis-eQTLs, that were associated with three genes *(BTN3A2, CHRNA5* and *GSTO2)*, were detected in whole blood (Supplementary Figure 12C). Of these, only *cis*-eQTLs associated with *BTN3A2* and *GSOT2* reached our significance threshold; eQTLs correlated with *CHRNA5* expression had a minimum pblood-Westra >7 x 10^−4^. Thus, whilst cis-eQTLs can be detected in whole blood in principle, approximately 90% did not reach genome-wide significance levels. This strongly suggests that eQTL analyses should either be performed in disease-relevant tissue wherever possible or that more extensive studies targeting minimally invasive tissues are necessary to identify suitable brain correlates.

## Discussion

In this study, we report the presence of eQTLs in brain tissue for SNPs that are associated with a risk of developing schizophrenia and/or affective disorders. Even with a stringent p-value cut-off of 5 x 10^−8^, we identified 1 288 cis-eQTLs and 297 trans-eQTLs that were correlated with expression of fifteen and twenty-four genes, respectively. These associations held across four brain regions from two independent studies (UKBEC and GTEx) of neurologically-normal individuals as well as through permutation analyses of 10 000 iterations, supporting the robustness of these findings.

Meta-analyses of eQTLs across different brain regions revealed diverging patterns between *cis* and *trans:* substantially more cis-eQTLs were significant when considering the four overlapping brain than trans-eQTLs. The pleiotropy of cis-eQTLs across different brain regions described here is consistent with their replicative behaviour previously reported. Considering all cis-eQTLs in UKBEC, 25-50% were also found in other eQTL datasets, even in non-synonymous tissues like human monocytes^29, 43^. In our study, ~35% of cis-eQTLs detected in brain were also present in GTEx whole blood samples, albeit at lower p-values. In contrast to *cis*, we could only detect trans-eQTLs for the same tissue regions between the two datasets. This is consistent with notion that trans-eQTLs may act in a tissue-dependent manner^44–46^. Given the differences in locations and associations of trans-eQTLs amongst the various brain regions, this supports the hypothesis that the biology governing these *trans*associations may be more influenced by the intracellular environment than the cis-eQTLs^17^.

Of the 1 288 cis-eQTLs detected, nearly two-thirds were located within introns or intergenic regions, consistent with previous studies^26, 41^. Meanwhile, nearly 2% were classified as causing missense mutations in the proteins encoded from genes harbouring the eQTL. The most significant eQTL that results in a missense mutation within the gene whose expression it is associated with, rs16969968, leads to an amino acid change of aspartic acid (for major allele *G)* to asparganine (for minor allele A) at position 398 in *CHRNA5*, a subunit in nicotinic acetylcholine receptors. This causes a change from a negatively charged side chain to one that is polar but uncharged. Interestingly, functional *in vitro* studies have demonstrated that receptors containing this missense mutation are less responsive to a nicotinic agonist than ones with the more common variant^34, 47^ This leads to reduced cell-depolarization and cholinergic signalling and is consistent with the reported hypofunction of cholinergic signalling in schizophrenia^48^. Moreover, pharmaceutical modulation of cholinergic signalling may improve outcomes for schizophrenia^49^.

There is already modest evidence that this and other eQTLs affecting *CHRNA5* expression are associated with schizophrenia and affective disorders^4, 50, 51^. However, rs 16969968 has also been shown to be associated with increased tobacco use^34, 52^ and incidence of lung cancer^53, 54^. Therefore, given the disproportionate percentage of individuals with mental illness that smoke^55^, further studies are needed to ensure that these eQTLs are not associated with a confounding behaviour in such individuals. Nonetheless, these cis-eQTLs for *CHRNA5* demonstrate how genetic analyses can identify variants that may increase disease-risk while concurrently being potential therapeutic targets.

Our analysis has also replicated several other cis-eQTLs found in other studies. Previously, 27 brain eQTLs were identified in a meta-analysis of GWAS SNPs associated with five neuropsychiatric disorders (including the three studied here^9^) using cortical expression data from five separate studies^56^. We identified one of these overlapping with our study: rs4523957 associated with expression of *SRR* on chromosome 17p13.3. Though this was not the most significant eQTL for this gene, the association was still below the genome-wide threshold at p_over_ = 2.2 x 10^−9^, FDR = 3.6 x 10^−7^. *SRR* encodes for an enzyme that converts L-to D-serine, which has be found to be lower in the CSF of patients with schizophrenia^57^, supporting the role of glutamatergic neurotransmission in the biology of schizophrenia and affective disorders^58, 59^ Indeed, it is also a candidate drug target for schizophrenia, again highlighting the potential for eQTL studies to identify pathobiology that might be targeted pharmacologically. *ZSCAN31*, also known as *ZNF323*, is another cis-eQTL associated gene that has been previously identified as significantly associated with schizophrenia, bipolar disorder and psychosis in both a GWAS^60^ and an eQTL study using 193 human pre-frontal cortex samples^61^. In our study, multiple cis-eQTLs associated with this gene were significant (p_over_ = 2.9 x 10^−8^ for the top eQTL). Likewise, two genes that were prioritized as putatively causal from integrative analysis of PGC-SCZ2 GWAS with both whole blood and UKBEC data^62^, *SNX19* and *NMRAL1*, were associated with cis-eQTLs with some evidence of significance, pover = 8.3 x 10^−5^ and 2.2 x 10^−6^, respectively.

Despite some overlap with previous studies, several reported associations were not significant here. *CACNA1C* and *ZNF804A* are two genes that have been implicated in schizophrenia and affective disorders through multiple studies, including GWAS and case/control brain expression analyses^63–67^. While both UKBEC and GTEx data contain expression information regarding these genes and the set of analysis-SNPs contain variants that are in *cis*, we did not find any significant eQTLs in our meta-analysis of UKBEC and GTEx across the four overlapping brain regions. For *CACNA1C* (minimum p_over_ = 0.45), the effect size estimates of the *cis*-eQTLs had different signs across the various regions (Supplementary Table 11). For *ZNF804A*, no detected eQTLs overlapped between all four regions (Supplementary Table 12). Further, of those that were in moderate-high LD with each other, the effect size estimates also demonstrated different signs. This suggests that some disease-implicated *cis*-eQTLs may have varying effects in different brain regions. These are not captured in this study but warrants further investigation.

The greatest number of trans-eQTLs detected in this study was predominantly located within the *MHC* locus on chromosome 6p21. eQTL analysis of peripheral blood and immune cells suggest that this region is enriched for trans-eQTLs^68, 69^, especially when restricting variants to those that are trait/disease-associated^70^, as was done in this study. However, given the poor replicability of trans-eQTLs thus far, we are cautious about over-interpreting these results; substantially larger sample sizes are needed to detect and validate such eQTLs. Nonetheless, the identification of tissue-specific trans-eQTLs that withstand permutation analysis gives confidence that our findings may be biologically significant.

Progress in understanding the biology governing schizophrenia and affective disorders has been hampered by difficulty in accessing disease-relevant tissue. Therefore, peripheral blood, as it is more accessible, has previously been used as a proxy for studying eQTLs of complex diseases^16, 42, 62^ However, it is unclear as to how robust findings from blood samples are with respect to studying diseases outside of the hematopoietic system. By comparing cis-eQTL results from brain and blood, we find that a minority of the brain-eQTLs were detected in blood and their significance (as marked by p-value) was greatly diminished. Therefore, studying eQTLs in disease-related tissues, and in this case brain tissue, may prioritize eQTLs for further validation and mechanistic studies. Indeed this has previously been demonstrated in prostate tissue for eQTLs associated with prostate cancer^71^.

In summary, we have identified robust *cis*- and trans-eQTLs associated with schizophrenia and affective disorders in human brain tissue. Of the eQTL-associated genes, many have been implicated previously, such as *CHRNA5*, while others are novel associations (i.e. *ZNF603P*) that now merit further analyses. We also demonstrate that eQTL analysis in disease-related tissues allows for prioritization of associations for follow-up studies in diseased-tissue. These results provide insight into putative mechanisms related to development of schizophrenia and affective disorders, thereby identifying potentially new therapeutic targets.

## Files

*Supplementary_Figures.pdf*

This file contains Supplementary Figures 1-12.

*Supplemenatary_Tables.xlsx*

This file contains Supplementary Tables 1-12, with each table on a separate sheet. *UKBEC_Affiliations.pdf*

This file contains the UK Brain Expression Consortium authors and respective affiliations.

## Acknowledgements

We thank the individuals and their support groups who have contributed to these studies and Sean Byars for help and advice with data analysis.

## Author Contributions

OGB, MI and CRS designed the study. OGB, APN, and CRS carried out analyses. OGB wrote the manuscript with assistance from APN, MI and CRS.

